# Estimation of the Acetylcholinesterase activity of honey bees in Nepal

**DOI:** 10.1101/2020.12.31.424928

**Authors:** Shishir Pandey, Shankar Gotame, Sachin Sejuwal, Basant Giri, Susma Giri

## Abstract

Decline in honey bee colonies possess a serious threat to biodiversity and agriculture. Prior detection of the stresses with the help of biomarkers and their management ensures honey bee’s survivability. Acetylcholinesterase (AChE) is a promising biomarker to monitor exposure of honey bees towards environmental pollutants. In this preliminary study, we measured AChE activity in forager honey bees collected from six districts of Nepal, Kathmandu, Bhaktapur, Lalitpur, Chitwan, Rupandehi and Pyuthan during autumn and winter seasons. We estimated AChE tissue and specific activities from bee’s heads using commercial kit based on Ellman assay and protein concentration using Lowry assay. In total, we collected 716 foragers belonging to *A. cerana, A. mellifera* and *A. dorsata*. A significant increase in all three parameters measured: AChE tissue activity, AChE specific activity and protein concentration was observed in winter samples. Both AChE tissue and specific activities were lower in *A. mellifera* compared to either *A. cerana* or *A. dorsata*. Protein concentration was higher in *A. mellifera* than in *A. dorsata* and lower than in *A. cerana*. We show correlation between both AChE tissue and specific activities and protein concentration across season and species and discuss possible factors contributing to the observations. Our results clearly indicate the presence of stress in the winter which is manifested through overexpression of the AChE. We recommend a detailed study to determine the factors accountable for the stresses for better management of honey bees in Nepal.

## Introduction

Pollination of the wild and cultivated crops by the bees is an important aspect of the ecological sustenance. Two-thirds of the food crops that make up the top 90% of each country’s national per capita supply of food plants (based on the analysis of FAO food supply data of 146 countries by Prescott-Allen & Prescott-Allen 1990) require animal pollinators where 73% of the crops species requiring pollinators rely to some extent on a variety of bees (Buchmann & Nabhan 1997). Although, recently the importance of native wild bee pollinators has been realized (e.g. Ollerton *et al.* 2012), the role of honey bees in crop pollination still remains important particularly in its native regions as well as in its non-native regions (Morse & Calderone 2000; Hein 2009; Hanley *et al.* 2015; Hung *et al.* 2018; as indicated by the transport of honey bees across locations for pollination services, Jones Ritten *et al.* 2018).

However, a growing number of investigations have reported a high rate of bee morbidity and mortality throughout the world, raising tremendous concerns over food security and maintenance of natural biodiversity (Ellis *et al.* 2010; Potts *et al.* 2010; Goulson *et al.* 2015). Multiple biotic and abiotic causative agents such as the infestation of parasitic mites, bacterial and viral infections, pesticide contamination, frequent queen losses, poor nutrition, genetic weakening, habitat fragmentations, cultivation of transgenic plants, migratory stress and climate change have been identified for driving the widespread decline in honey bee populations (reviewed in Potts *et al.* 2010; vanEngelsdorp & Meixner 2010; Vanbergen & Initiative 2013). Some of these environmental stressors have also been implicated in the episodes of colony collapse disorder (CCD), an inexplicable disappearance that resulted in the sharp decline of worker honey bees (Cox-Foster *et al.* 2007). The significant drop in honey bee populations due to agriculture intensification, pervasive use of agriculturally administered insecticides and environmental pollution has drawn much attention.

One group of insecticide, neonicotinoids, which act as the agonist of neurotransmitter acetylcholine (ACh, Matsuda *et al.* 2001; Tan *et al.* 2007) are found to have lethal impacts on honey bees (reviewed in Lundin *et al.* 2015; Benuszak *et al.* 2017). Studies have made clear that the exposure of bees to sublethal level of insecticides lead to disrupted cholinergic neurotransmission resulting in the impairment of all cognitive functions such as impairment of learning and memory, disrupted navigation, decline in immunity and foraging activity among others (Palmer *et al.* 2013; Tan *et al.* 2015; Chmiel *et al.* 2020). In addition, adverse effects on queen bee fecundity (Wu-Smart & Spivak 2016) and genetic diversity of the colony (Forfert *et al.* 2017) have also been reported. Trophallactic transfer of pesticide contaminated nectar and pollen to the colony by the forager bees results in the contamination of bee products like honey, royal-jelly, propolis and wax. Detection of insecticides residues in honey has been reported in a worldwide survey which has negative human health consequences as well (Mitchell *et al.* 2017).

Different environmental stressors can act alone or synergistically act together to exacerbate the health of honey bees (e.g. Johnson *et al.* 2013; Straub *et al.* 2019). Identification of suitable biomarkers for the prior detection of stress in honey bee colonies, therefore, becomes crucial for proactive management (Kim *et al.* 2019). Various biomolecules including glutathione-S-transferase, acetylcholinesterase (AChE), metallothioneins, alkaline phosphatase, nicotinic acetylcholine receptors (nAChRs), cytochrome P450 oxidase, vitellogenin and α-tocopherol are used to assess the effects of diverse stressors in honey bees (Badiou-Bénéteau *et al.* 2013; Christen & Fent 2017; Gauthier *et al.* 2018). Enzymatic biomarker, AChE is targeted as a promising candidate, because of its vulnerability to express differentially in relation to the pesticide exposure (Boily *et al.* 2013; but see Kim *et al.* 2019), seasonal condition (Kim *et al.* 2017), landscape (Badiou-Bénéteau *et al.* 2013) and nutrition status (Kim *et al.* 2017, 2019). AChE rapidly hydrolyses ACh at the cholinergic synapses, thus allowing precise control and modulation of the neural transmission (Badiou *et al.* 2008). Organophosphate and carbamate classes of insecticides are known to exert an inhibitory effect on AChE (Belzunces *et al.* 2012; Palmer *et al.* 2013; Williamson *et al.* 2013). Studies with neonicotinoids at sublethal doses have shown upregulated (Boily *et al.* 2013), downregulated (Li *et al.* 2017) and unchanged (Alburaki *et al.* 2017; Gauthier *et al.* 2018) AChE activity in honey bees under laboratory and semi-field conditions.

Beekeeping is an important source of income generation in Nepal. Out of five species of honey bees found, four of them are indigenous, *viz. Apis florea* F., *Apis cerana* F., *Apis dorsata* F., and *Apis laboriosa* S. (Allen 1995). Commercial beekeeping started after the introduction of *Apis mellifera* L. during 1990 (Aryal *et al.* 2015). Inadequate intervention in colony management, mite infestation and brood diseases, among other factors, are reported for the decline in honey bee colonies in Nepal (Thapa *et al.* 2018). Given that the modern unsustainable agricultural techniques and subsistence climate change act insidiously, the inevitable threat in the extinction of native Himalayan species of honey bees is alarming. Therefore, the status of biomarkers that are intricate to the honey bees’ survivability needs to be addressed as a pre-emptive strategy in Nepal. This is a preliminary study on the estimation of AChE activity from forager honey bees collected from different anthropogenic sites of Nepal. The determined AChE activity herein, can serve as an index that can be used as a reference for further detailed studies.

## Materials and Methods

### Sample collection

We collected female worker honey bees (N = 716) from various locations in Nepal using the net swipe method (Fig 1). Sample collection was done in two phases. In the first phase, bees were collected in 2019 during November and first week of December when the bees were still foraging and were assigned as autumn samples. We collected a total 427 *A. cerana* from Kathmandu Valley (hilly region) and 90 *A. mellifera* and 24 *A. dorsata* from Chitwan (low-lying Terai region).

**Fig 1.**
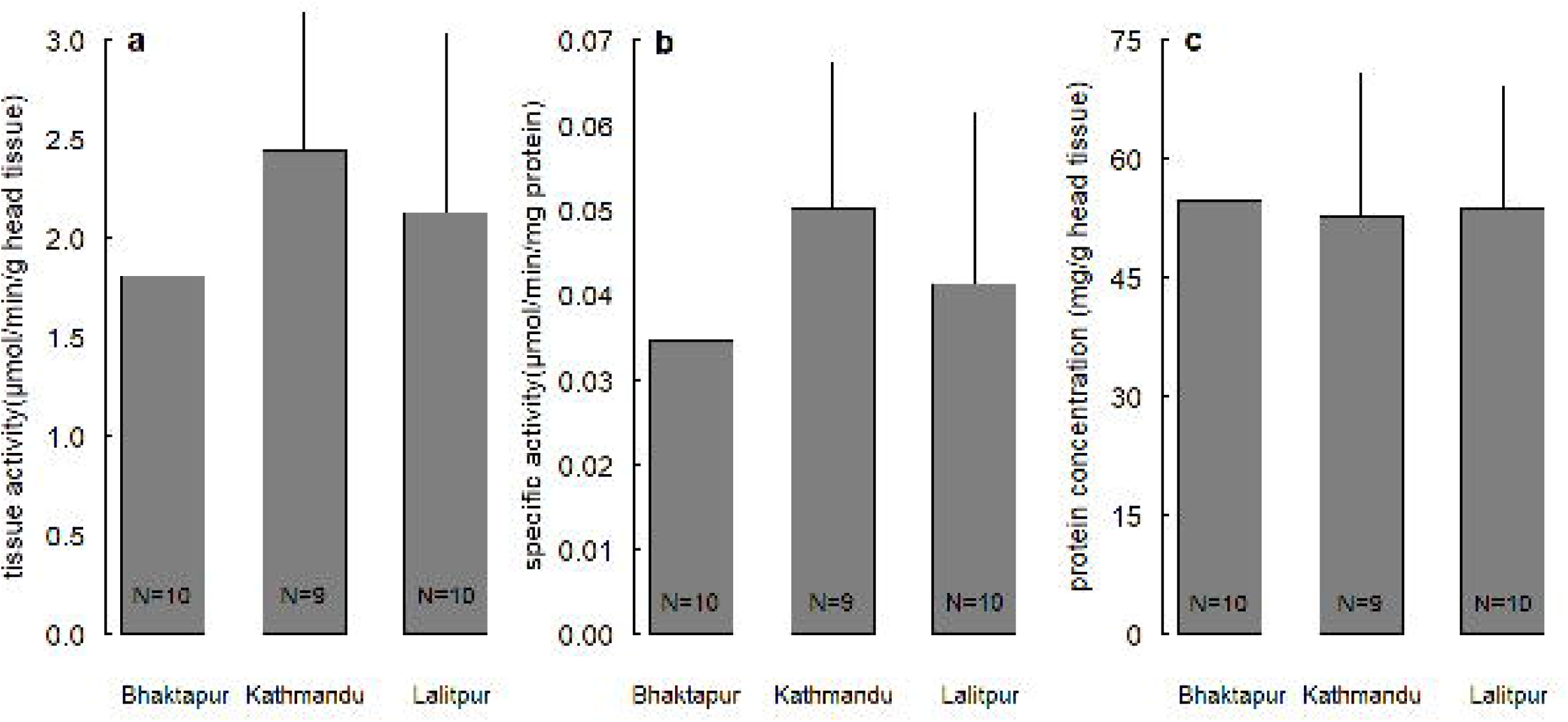
Map of Nepal with sample collection locations (a). Partial map of Nepal showing sample collection regions and seasons (autumn/winter) (b).

In the second phase, bees were collected in early spring (February, 2020) directly from bee hives (N = 175) and were assigned as winter samples. We collected a total of 115 *A. cerana* and 60 *A. mellifera* from six apiaries located in Rupandehi (low-lying Terai region), Kathmandu and Pyuthan (hilly regions, see Table 1 for details). All the collected bee samples were stored in a freezer at the hotel and transported to the lab on ice box and were kept at −20 °C until further analyses (usually within 10 days). Bees were identified using personal knowledge/communication and some guides including Michener *et al.* (1994) and Walker (2006).

**Table 1:**
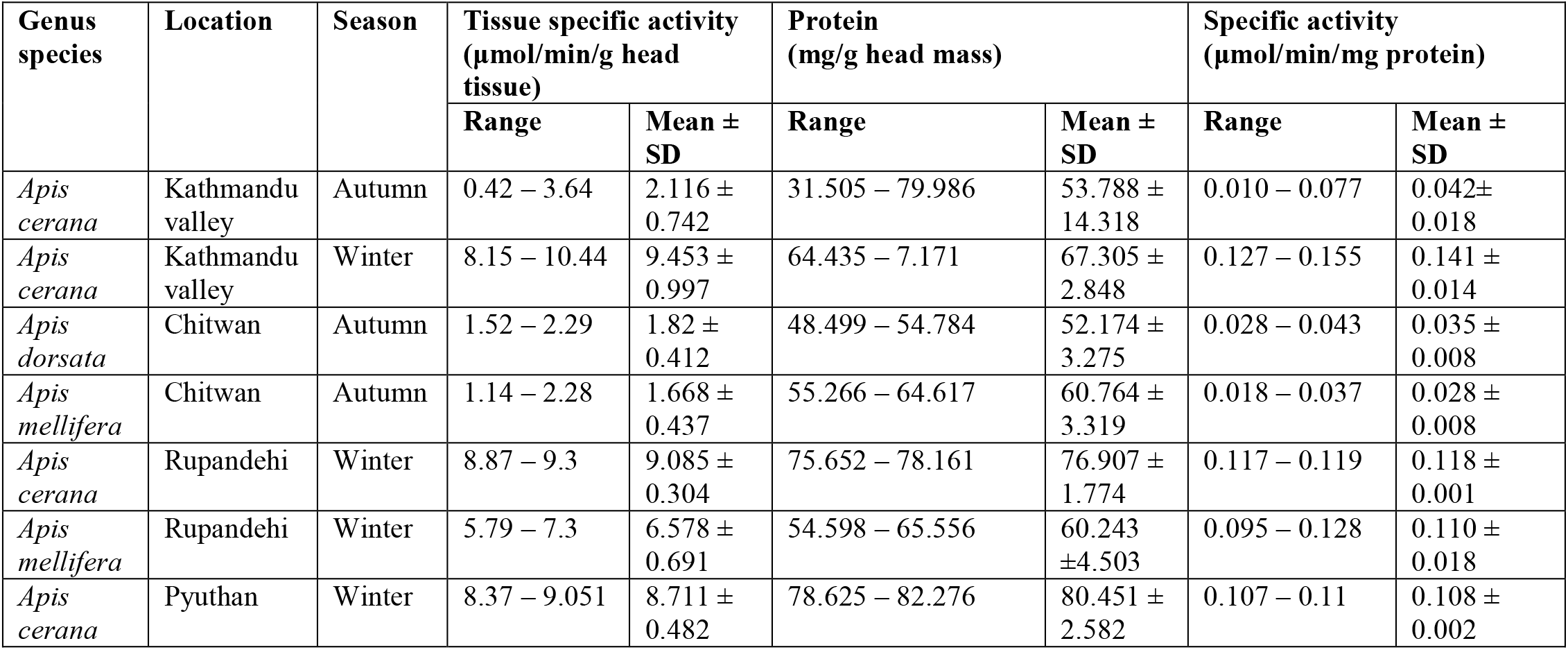
Mean ± SD values for all three parameters measured in honey bees across season and species.

| Genus species | Location | Season | Tissue specific activity ($\mu\text{mol}/\text{min}/\text{g}$ head tissue) | | Protein ($\text{mg}/\text{g}$ head mass) | | Specific activity ($\mu\text{mol}/\text{min}/\text{mg}$ protein) | |
| --- | --- | --- | --- | --- | --- | --- | --- | --- |
| | | | Range | Mean ± SD | Range | Mean ± SD | Range | Mean ± SD |
| *Apis cerana* | Kathmandu valley | Autumn | 0.42 – 3.64 | 2.116 ± 0.742 | 31.505 – 79.986 | 53.788 ± 14.318 | 0.010 – 0.077 | 0.042 ± 0.018 |
| *Apis cerana* | Kathmandu valley | Winter | 8.15 – 10.44 | 9.453 ± 0.997 | 64.435 – 7.171 | 67.305 ± 2.848 | 0.127 – 0.155 | 0.141 ± 0.014 |
| *Apis dorsata* | Chitwan | Autumn | 1.52 – 2.29 | 1.82 ± 0.412 | 48.499 – 54.784 | 52.174 ± 3.275 | 0.028 – 0.043 | 0.035 ± 0.008 |
| *Apis mellifera* | Chitwan | Autumn | 1.14 – 2.28 | 1.668 ± 0.437 | 55.266 – 64.617 | 60.764 ± 3.319 | 0.018 – 0.037 | 0.028 ± 0.008 |
| *Apis cerana* | Rupandehi | Winter | 8.87 – 9.3 | 9.085 ± 0.304 | 75.652 – 78.161 | 76.907 ± 1.774 | 0.117 – 0.119 | 0.118 ± 0.001 |
| *Apis mellifera* | Rupandehi | Winter | 5.79 – 7.3 | 6.578 ± 0.691 | 54.598 – 65.556 | 60.243 ± 4.503 | 0.095 – 0.128 | 0.110 ± 0.018 |
| *Apis cerana* | Pyuthan | Winter | 8.37 – 9.051 | 8.711 ± 0.482 | 78.625 – 82.276 | 80.451 ± 2.582 | 0.107 – 0.11 | 0.108 ± 0.002 |

### Sample Preparation

To increase the volume of sample, we pooled 8-15 bees together from each sample collection site for all of the tests performed. Any water or nectar droplet present on the surface of bees were soaked with a clean paper towel. We then cryo-cleaved heads from the body using sterilized scalpel and weighed individually in the balance (AP135W, Shimadzu, Japan).

We next transferred ~100 mg (range = 62 to 170 mg) of tissues to a glass Teflon homogenizer and ground using appropriate volume of freshly prepared low salt triton (LST) buffer containing 1% (v/v) TritonX-100 (MB031, HIMEDIA^®□^, India), 10 mM NaCl and 15mM sodium phosphate dibasic, pH 7.5 to make a 10% (w/v) extract as suggested by Boily *et al.* (2013). We used ice cubes to maintain colder temperatures during homogenization. The homogenate was then transferred to 2 mL centrifuge tubes, homogenizer rinsed with additional 200 μL LST and recovered into the same tube and then centrifuged at 17,000 x g for 30 min at 4 °C (NF 800R, Nüve, Turkey). Supernatants were pooled carefully while the pellets were subjected to second homogenization with 500 μL LST and centrifuged as above. Supernatants pooled out were combined with previous and stored at −20 °C for subsequent analyses.

### Acetylcholinesterase and protein assay

We determined the acetylcholinesterase activity of each sample following Ellman method (Ellman *et al.* 1961) using a commercial KIT (MAK 119, Sigma-Aldrich, USA). Briefly, the Ellman reaction involves the hydrolysis of acetylthiocholine by AChE to produce thiocholine and acetate. Thiocholine, in presence of dithiobisnitrobenzoate, is reduced to the yellow anion of 5-thio-2-nitrobenzoic acid imparting yellow color to solution, the intensity of which is then compared to the standard for quantification of AChE activity. We measured the intensity of yellow color developed from the Ellman reaction in a multiplate reader (BioTek, USA) at 412 nm. The absorbance read at 10 min was subtracted from the absorbance read at 2 min to obtain AChE activity. All readings were corrected from the blank reaction. Sample AChE activity in μmol/min/L was then calculated by comparing with the calibrator colour. One unit of AChE is the amount of enzyme that catalyses the production of 1 μmol of thiocholine per min at 25 °C at pH 7.5 (“Product information: Acetylcholinesterase Activity Assay Kit” 2013). AChE specific activity was expressed in μmol/min/mg protein, AChE tissue activity was expressed in μmol/min/mg tissue and protein concentration was expressed in mg/g head tissue.

We quantified protein in the sample following the Lowry method as described by Waterborg & Matthews (1994). Briefly, the peptide bonds of proteins react under alkaline conditions with Cu^2+^ to produce Cu^+^, which, along with the radical groups of amino acids then reacts with Folin reagent which is then reduced to molybdenum/tungsten blue. The intensity of the color is then compared with the standard calibration curve for quantification of protein in the sample. However, the use of non-ionic detergent TritonX-100, that is used to dissociate protein from membranes interfere with this method of protein determination by producing precipitate upon addition of the Folin reagent. We eliminated the precipitation of Folin reagent with the addition of 0.5% (w/v) sodium dodecyl sulfate in the alkali reagent as suggested by Dulley and Grieve, 1975. Bovine serum albumin (MB083, HIMEDIA^®□^, India) was used as a standard for protein estimation. Protein concentration was reported as mg of protein per g of head tissue.

### Statistical analyses

We report AChE activities (tissue and specific activities) and protein concentrations in bees from varying locations, season and species as mean ± standard deviation (SD). The number of pools containing 8-15 honey bees ranged from N = 2 to 10 according to the criteria studied. Given smaller sample size (N < 10), we performed variance (package “stats”), coefficient of variation (package “goeveg”) and ANOVA tests as necessary to check if data for a given species and season can be combined for varying location with similar conditions. Normality of the data was tested using Shapiro-Wilk normality test and difference between coefficients of variation between two groups was tested using R package “cvequality” (Marwick & Krishnamoorthy 2019). Owing to the normal distribution of data, we then performed Welch Two Sample t-test to compare between two variables and one-way analysis of variance (ANOVA) to compare among more than two variables followed by the contrast procedure with Tukey’s honestly significant difference (HSD) test. Pearson product-moment correlation was used to evaluate the correlation between AChE tissue activity and protein concentration. Statistical analyses were performed in R language using RStudio (version 1.2.5033) (R Core Team, 2020).

## Results

We report AChE tissue activity (μmol/min/mg head tissue), AChE specific activity (μmol/min/mg protein) and protein concetration (mg/g head tissue) for a pool (N) of 50 constituting a total (n) of 716 bees representing different locations, seasons and species. Coefficient of variation was higher for AChE tissue activity for some combinations and for AChE specific activity for others (data not shown). As such, we show statistical tests for both parameters.

Given similar geographic, climatic and urbanization conditions as well as located within the same valley, we checked if data for within Kathmandu Valley (i.e. from Bhaktapur, Kathmandu and Lalitpur districts) for a given species and season can be combined for further comparisons. Although these districts showed significant difference in coefficient of variation for all three parameters measured (all *P* < 0.066), we found no significant difference between groups (AChE tissue activity: F_2, 26_ = 1.84, *P* = 0.178; AChE specific activity: F_2, 26_ = 1.90, *P* = 0.17; protein concentration: F_2, 26_ = 0.05, *P* = 0.955). We, therefore, combined the data for a given species and season for further comparisons (Table 2; Fig 2 shows values separately for these districts). The basic values (range and mean ± SD) for all groups and the parameters measured are shown in Table 3.

**Fig 2.**
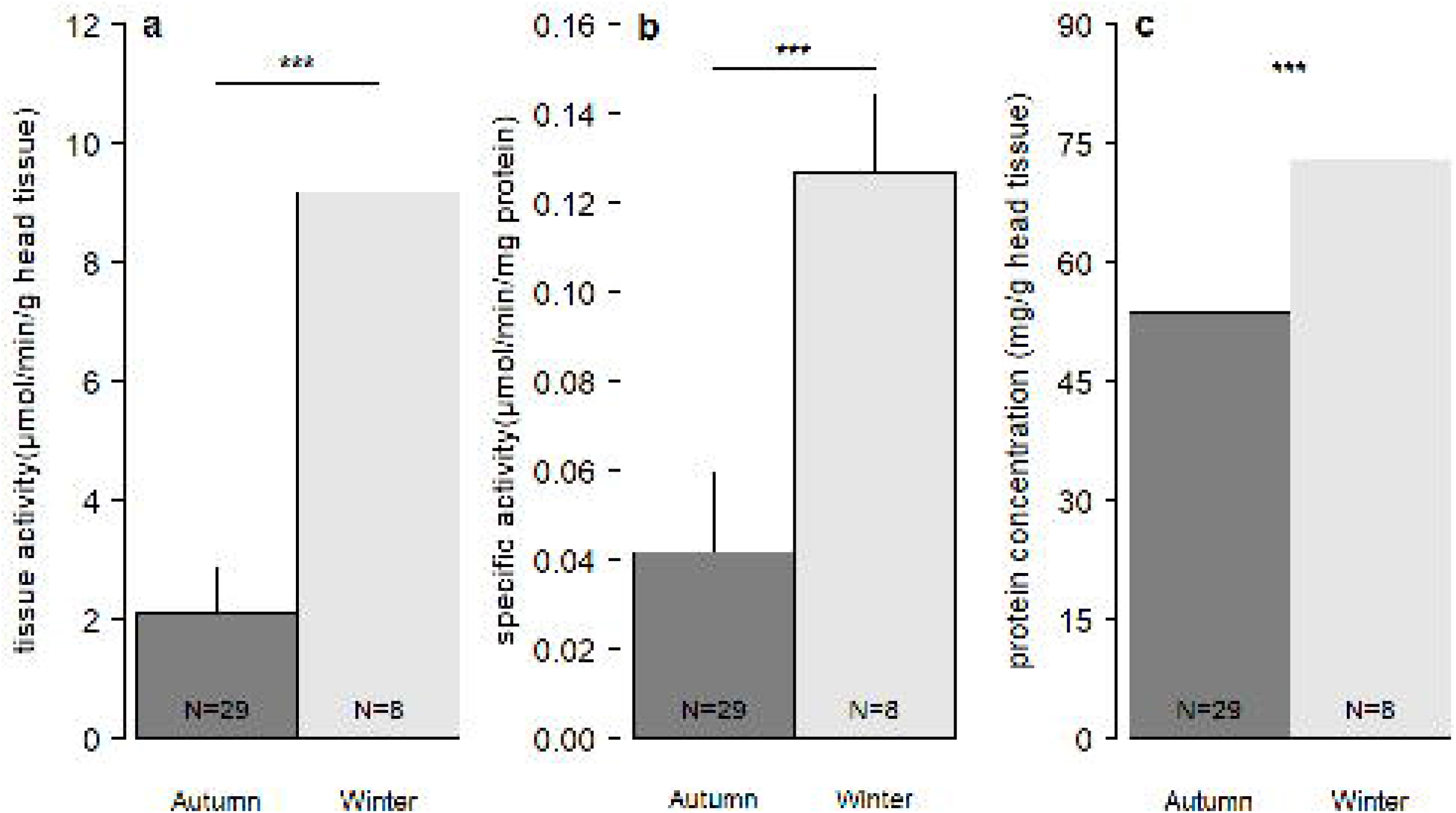
Bar chart showing AChE tissue activity (a), AChE specific activity (b) and protein concentration (c) in Autumn *A. cerana* collected from Kathmandu, Bhaktapur and Lalipur.

**Table 2:** Details of sample collection.

| Genus species | District | Season | # of pools | # of bees | # per pool |
| --- | --- | --- | --- | --- | --- |
| *Apis cerana* | Kathmandu | Autumn | 9 | 135 | 15 |
| *Apis cerana* | Bhaktapur | Autumn | 10 | 144 | 11-15 |
| *Apis cerana* | Lalitpur | Autumn | 10 | 148 | 13-15 |
| *Apis cerana* | Kathmandu | Winter | 4 | 60 | 15 |
| *Apis dorsata* | Chitwan | Autumn | 3 | 24 | 8 |
| *Apis mellifera* | Chitwan | Autumn | 6 | 90 | 15 |
| *Apis cerana* | Rupandehi | Winter | 2 | 30 | 15 |
| *Apis mellifera* | Rupandehi | Winter | 4 | 60 | 15 |
| *Apis cerana* | Pyuthan | Winter | 2 | 25 | 12-13 |

### Seasonal variation in AChE activity and protein concentration

We compared AChE specific activity and protein concentration between *A. cerana* collected during autumn and winter. A total of 29 pools (n = 427, see Table 3) comprising combined autumn data for Kathmandu Valley (Bhaktapur, Kathmandu and Lalitpur) were compared with 8 pools (n = 115) comprising data for Kathmandu, Pyuthan and Rupandehi districts. Given low sample size when only compared for *A. cerana* for individual location, we evaluated the suitability of combining all *A. cerana* samples collected during the winter season from these three districts. An asymptotic test for the equality of coefficients of variation showed no significant differences (*P* = 1) as well as the CV was not that high (14%). Hence, we combined data for *A.cerana* collected from Kathmandu (N = 4), Pyuthan (N = 2) and Rupandehi (N = 2) in winter for comparison with autumn data.

All three parameters, AChE tissue activity (winter: 9.175 ± 0.762 μmol/min/g head tissue, autumn: 2.116 ± 0.742 μmol/min/g head tissue; t_11_ = −23.34, *P* < 0.001), AChE specific activity (winter: 0.127 ± 0.018 μmol/min/mg protein; autumn: 0.042 ± 0.018 μmol/min/mg protein; t_11_ = −11.96, *P* < 0.001) and protein concentration (winter: 72.992 ± 6.606 mg/g head tissue; autumn: 53.788 ± 14.318 mg/g head tissue; t_26_ = −5.43, *P* < 0.001) were significantly higher in winter compared to that in autumn (Fig 3). Though not significant, AChE tissue activity showed a positive correlation with protein concentration in autumn (*r* = 0.21, *P* = 0.271) and a negative correlation in winter (*r* = - 0.28, *P* = 0.510). Whereas, a significant negative correlation was established between AChE specific activity and protein concentration in both autumn (*r* = −0.53, *P* = 0.003) and winter (*r* = - 0.80, *P* = 0.017) *A. cerana* samples.

**Fig 3.**
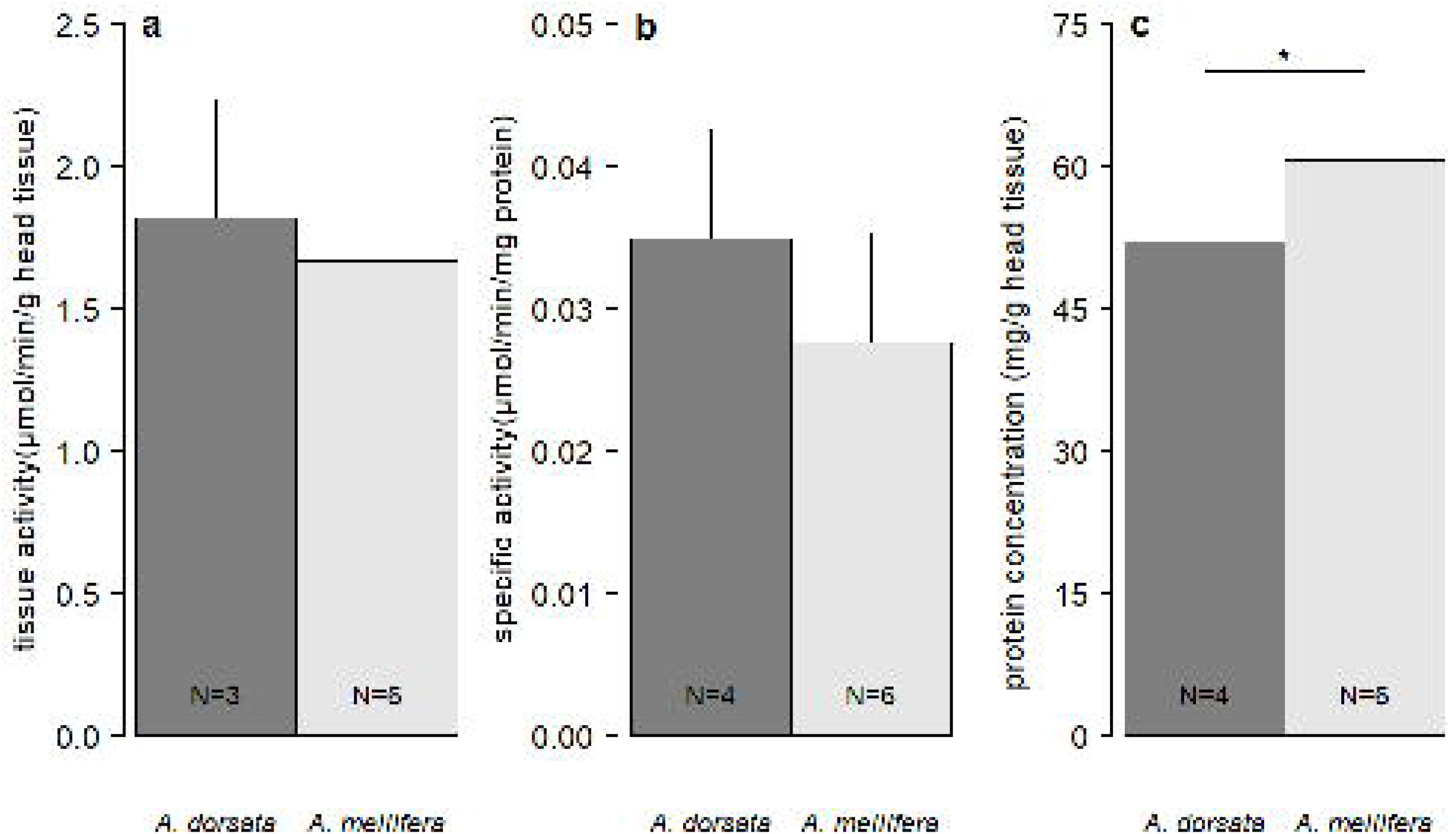
Seasonal comparison of AChE tissue activity (a), AChE specific activity (b) and protein concentration (c) in *A. cerana* samples.

### Species-wise variation in AChE activity and protein concentration

We compared AChE specific activity and protein concentration among among *A. dorsata* (N = 3) and *A. mellifera* (N = 6) collected from Chitwan during autumn. We also compared the parameters among *A. cerana* (N = 8) and *A. mellifera* (N = 4) collected during winter. Data for *A. cerana* from Kathmandu, Rupandehi and Pyuthan were combined after asymptotic and CV tests as explained above.

We found no significant difference in AChE tissue activity between *A. dorsata* (1.820 ± 0.412 μmol/min/g head mass) and *A. mellifera* (1.668 ± 0.437 μmol/min/g head mass) collected from Chitwan (t_4_ = 0.5, *P* = 0.635). AChE specific activity was also not different between species (*A. dorsata:* 0.035 ± 0.008 μmol/min/mg protein; *A. mellifera:* 0.028 ± 0.008 μmol/min/mg protein; t_4_ = 1.4, *P* = 0.242). Protein concentration was, however, significantly higher in *A. mellifera* (60.764 ± 3.319 mg/g head tissue; t_4_= −3.7, *P* = 0.02; *A. dorsata:* 52.174 ± 3.275 mg/g head tissue; Fig 4). A weak, yet positive correlation was found between AChE tissue activity and protein concentration in *A. dorsata* (*r* = 0.13, *P* = 0.919) but a negative correlation was found between these parameters in *A. mellifera* (*r* = −0.25, *P* = 0.634). AChE specific activity, on the other hand, showed negative correlation with protein concentration in *A. dorsata* (*r* = −0.13, *P* = 0.919) and a negative correlation in *A. mellifera* (*r* = −0.44, *P* = 0.382).

**Fig 4.**
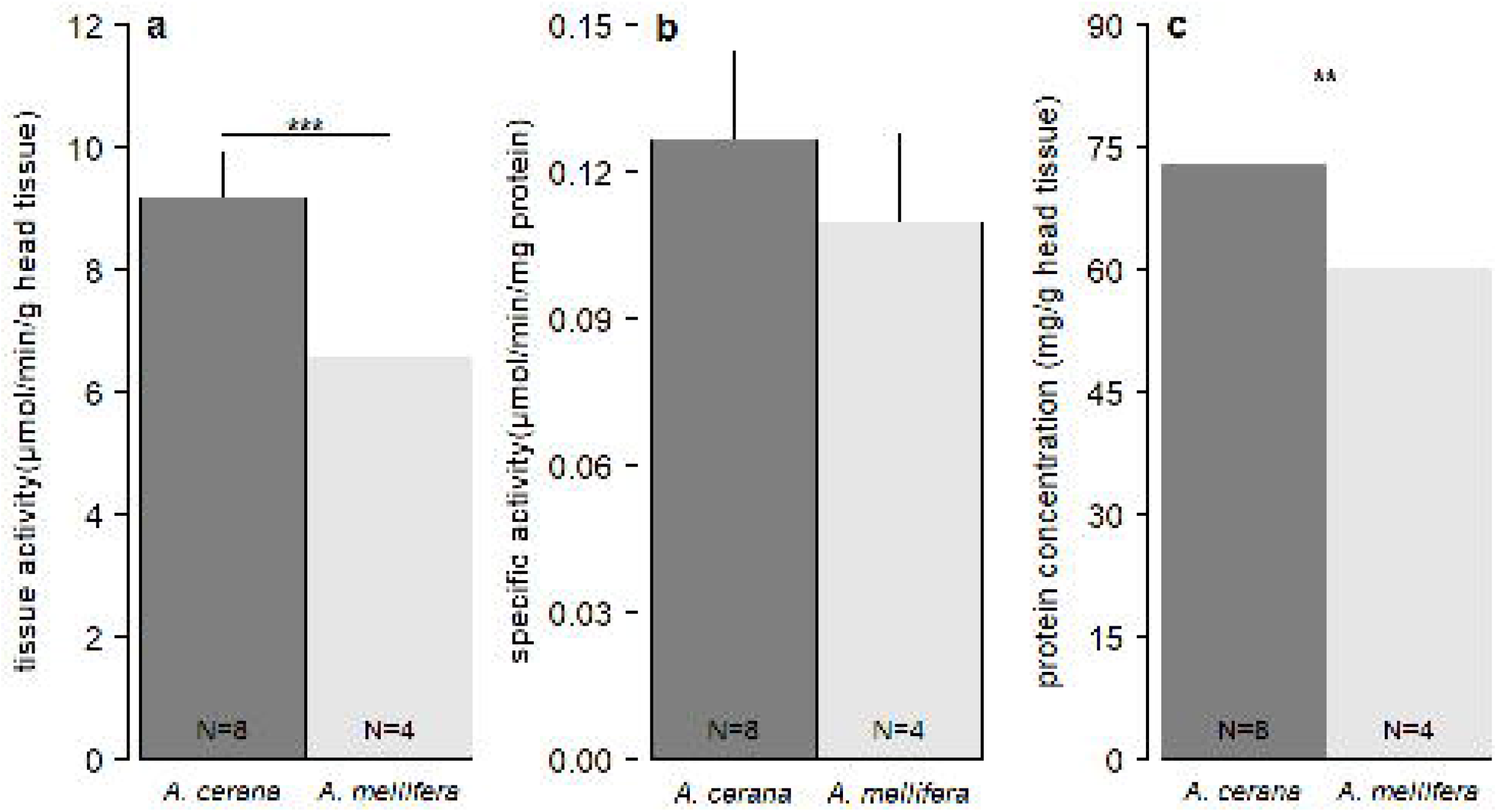
Species-wise comparison of AChE tissue activity (a), AChE specific activity (b) and protein concentration (c) in honey bees collected from Chitwan

AChE tissue activity and protein concentration were significantly higher in *A.cerana* (9.175 ± 0.762 μmol/min/g head mass and 72.992 ± 6.606 mg/g head tissue respectively) than in *A. mellifera* (6.578 ± 0.691 μmol/min/g head mass and 60.243 ± 4.503 mg/g head tissue; t_7_ = 5.9, *P* < 0.001 and t_9_ = 3.9, *P* = 0.004 respectively). AChE specific activity, on the other hand, didn’t differ significantly between species (t_6_ = 1.6, *P* = 0.167; Fig 5). As already mentioned above, a negative correlation was established between protein concentration and AChE tissue activity in *A. cerana* (*r* = - 0.28, *P* = 0.510) as well as in *A. mellifera* (*r* = - 0.54, *P* = 0.458). Likewise, a significant negative correlation was established between protein concentration and AChE specific activity in *A. cerana* (*r* = - 0.80, *P* = 0.017) and not significant in *A. mellifera* (*r* = - 0.83, *P* = 0.168).

**Fig 5.**
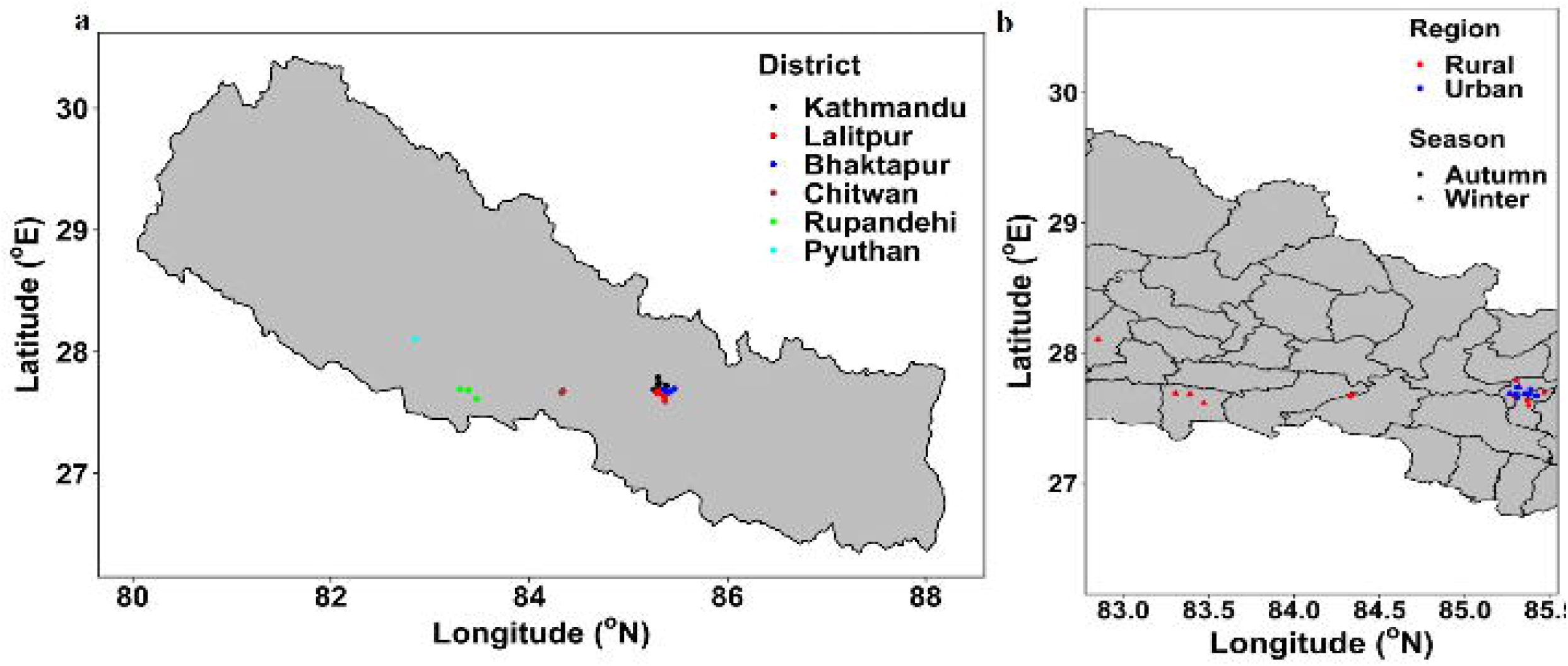
Species-wise comparison of AChE tissue activity (a), AChE specific activity (b) and protein concentration (c) in honey bees collected from Rupandehi

## Discussion

In the current study, we estimated AChE activity in forager honey bees (*A. cerana, A. dorsata* and *A. mellifera*) collected from different anthropogenic sites during autumn and winter seasons in Nepal. Any manifestation in the biomarker level is pertinent with forager bees because they are directly encountered with xenobiotic pollutants during their foraging activity. Moreover, biological variations linked to age, gender and spawning period can be reduced by sampling foragers only (Badiou-Bénéteau *et al.* 2013). Honey bees are extremely sensitive to pesticide exposure than pest insects because of the diminished repertoire of the genes for innate immunity and detoxifying enzymes such as cytochrome P450 (Weinstock *et al.* 2006). Mortality rates related to various stress factors were comprehensively conducted in western honey bees but it is largely underexplored with eastern counterpart, even though more than 50% of the global economic value of insect pollination originates from Asia (Gallai *et al.* 2009; reviewed in Lundin *et al.* 2015).

Honey bee biomarkers related to antioxidation, detoxification, immunity and neurotoxicity are promising tools for monitoring environmental quality. AChE embodies a general biomarker for neurotoxic effects because not only does it show systematic response to pesticide but also to heavy metals, radioactive elements and complex mixtures of industrial and urban pollutants (Porrini *et al.* 2002; Frasco *et al.* 2005; Khalifa *et al.* 2020). Kathmandu Valley is one of the most polluted city in the world (Neupane *et al.* 2020). High altitude of ~1300 m in Kathmandu Valley favours the beekeeping of native *A. cerana* as it is better adopted in harsh mountain environments than high honey yielding *A. mellifera* (Pokhrel *et al.* 2006). Although, there were no significant differences in the AChE activity across districts of the Kathmandu Valley, low AChE activity seen in honey bees collected from Bhaktapur (Fig 2) could suggest the presence of more neurotoxic compounds in this region as we collected more honey bees from agricultural sites in Bhaktapur (*see* Table 1). A home to over 120 brick kilns (which lead to the emission of various air pollutants, e.g. Rajarathnam *et al.* 2014) and rapid development, the average air pollution level is higher in Bhaktapur than in Kathmandu (Shrestha 2016). With its increasing population (Government of Nepal, National Planning Secretariat 2014), the synergistic effects of agricultural and industrial pollutants, along with other anthropogenic stressors may explain the lower AChE levels in Bhaktapur (see Badiou-Bénéteau *et al.* 2013), although a detailed study is needed for confirmation.

Likewise, ecological and species-wise variations can also alter AChE activity. Chitwan district is a major beekeeping hotspot of Nepal. The climatic conditions and altitude (~415 m) favors the existence of three species of honey bees (*viz., A. mellifera, A. cerana* and *A. dorsata*). *A. mellifera* is mostly kept for the commercial production of honey. From fall to autumn, mustard is cultivated in this region which provides ample pollen and nectar for honey bees. Beekeepers place their beehives around the cultivation site and allow for foraging for around one to three months (Personal communication, 2019). However, this is also the peak period for the use of insecticides required for the protection of mustard against pests. Estimated pesticide amount of 151 g active ingredient (a.i.) per hectare of arable land was recorded in Nepal in 2008 and is on the increasing trend (see Atreya & Sitaula 2010). The situation is worsened when the administration of insecticides or pesticides is more than the required amount. Prevalence of such practise was reported leading to the AChE depression in the farm workers of Nepal (Atreya *et al.* 2012). During sample collection time, we observed the administration of cocktails that includes acetamiprid (second generation Chloro-neonicotinoids most widely used in Asia), chlorpyrifos (organophosphate) and cypermethrin (pyrethroid). In addition, studies indubitably address the presence of stressors in this region (e.g. Mahapatra *et al.* 2018). The pesticide level in honey from a hive provides the measure of the contamination of the surrounding landscape. In a worldwide survey, which includes Nepal, honey collected from beehives in Chitwan was found to be contaminated with neonicotinoids such as imidacloprid (0.047 ng/g) and thiamethoxam (0.026 ng/g) (Mitchell *et al.* 2017). The later research provides a stronghold for the possible threat to the honey bee colonies due to the pesticide. We observed higher specific activity in *A. cerana* samples collected during Autumn from Kathmandu Valley compared to that in *A. mellifera* collected during the same season from Chitwan (mean=0.042±0.018 versus 0.028±0.008, respectively; t_18_=-3.1, *P* = 0.006). Similarly, the value was significantly higher in winter *A. cerana* collected from Kathmandu Valley than from Rupandehi (mean = 0.141±0.014 versus 0.118±0.001 respectively, t_3_=3.11, *P* = 0.051). However, in order to establish the depression due to insecticide is challenging without a control group given multiple factors.

In the winter season, we observed a remarkable increase in both tissue and specific AChE activities in *A. cerana* samples. This clearly indicates the presence of stress in winter which is mediated through AChE enzyme. A recent report discreetly suggested the role of soluble AChE1 (acetylcholinesterase 1) in the stress response or stress management during the cold conditions. Although, AChE1 has significantly less catalytic activity, given that it is abundantly expressed during winter months, it can still augment to the hydrolysis of ACh to produce high AChE activity (Kim *et al.* 2012). Previous study also suggested the involvement of AChE in the defense against xenobiotic pollutants (Kim *et al.* 2012, 2017). Therefore, other than the cold condition itself, the level of environmental pollutants which is significantly higher during winter months could have also probably contributed to the increase in the AChE activity. Neonicotinoids act on nAChRs by mimicking ACh which in turn over stimulates neural transmission so as a natural response AChE is produced to end the hyperactive nerve transmission (Samson-Robert *et al.* 2015). Therefore, high AChE levels are sought as a bioindicator for the exposure to neonicotinoids (Boily *et al.* 2013). In our study, honey bees collected in winter were fed mostly with sugar syrup and their foraging activity is restricted due to the cold condition; it is less likely that the possibility of high AChE level is due to neonicotinoids exposure particularly from afar. However, high AChE level due to the neonicotinoids exposure cannot be neglected because bees wax being lipophilic in nature can have high bioaccumulation of pesticides (Calatayud-Vernich *et al.* 2019) and cold conditions in winter may render foragers to have an increased contact with the contaminated beehives possibly leading to higher AChE activity.

In this study, both tissue and specific AChE activities were lower in *A. mellifera* (whether significant or not) compared to either *A. cerana* or *A. dorsata* indicating that the latter two species are more sensitive to the stressors than *A. mellifera* as previously reported (*A. cerana* vs *A. mellifera*; e.g. Li *et al.* 2017). A difference in effects of pesticides (as measured by LD50) across species of honey bees previously reported, with possibilities for behavioural and/or physiological explanations. A few possible cause of the observed differences may be due to the differences in management regimes (managed vs wild or types of managements), repeated disturbance of hives by bee keepers or honey hunters, differences in the physiological mechanisms for the detoxification of pesticides (among various other stressors), or the location of hives (Lee *et al.* 2016).

Previously, the status of the protein concentration related to the stressful environment have often been overlooked. In our study, similar to the AChE activity, it was interesting to find significant increment in the protein concentration in the winter samples than in the autumn samples. A significant increase in protein concentration was found for *A. mellifera* compared to *A. dorsata* but lower than in *A. cerana*. The relationship between AChE tissue/specific activity and protein concentration was rather complicated and varied with the season, species and location. However, it can be noted from our results that when significant, an increase in AChE activity is associated with increased protein synthesis whether across season or across species although opposite was observed for *A. mellifera* collected from Chitwan (*see* Fig. 3, 4 & 5). Increase in the signalling pathways associated with the stress, among others, could have repercussions in the synthesis of large amounts of proteins (Scharlaken *et al.* 2007), which may explain the results obtained in this study.

In summary, from this preliminary report, it is apparent that the cellular response is manifested through the AChE enzyme, which, can be used as an effective biotool for monitoring seasonal stresses in honey bees. However, in this study we were unable to pinpoint a distinct or synergistic factor that is accountable for the overexpression of the AChE enzyme. Pesticide residue analysis, and study of disease in the winter beehives along with lab-based experiments can narrow down some contributable factors. Using AChE as a suitable biomarker, a comprehensive study incorporating a large sample size with longitudinally collected samples across all seasons and multiple years are needed in Nepal.

## Ethical consent

National guidelines for the use of animals were followed.

## Acknowledgement

We sincerely thank Prajwal Rajbhandari and Mitesh Shrestha from Research Institute for Bioscience and Biotechnology (RIBB), Lalitpur, Nepal, for providing cooling centrifuge. We thank Jhalak Paudel (Research Intern, KIAS) for helping with identification of bee samples.

## References

Alburaki, M., Steckel, S.J., Chen, D., McDermott, E., Weiss, M., Skinner, J.A., et al. (2017). Landscape and pesticide effects on honey bees: forager survival and expression of acetylcholinesterase and brain oxidative genes. Apidologie, 48, 556–571.

Allen, M.F. (1995). Bees and beekeeping in Nepal. Bee World, v. 76(4) p.

Aryal, S., Thapa, R., Wongsiri, S., Lee, M.L., Choi, Y.-S., Ahn, Y.-J., et al. (2015). An overview of Beekeeping Economy and Its Constraints in Nepal. J. Apic., 30, 135–142.

Atreya, K. & Sitaula, B.K. (2010). Mancozeb: growing risk for agricultural communities? Himal. J. Sci., 6, 9–10.

Atreya, K., Sitaula, B.K., Overgaard, H., Bajracharya, R.M. & Sharma, S. (2012). Knowledge, attitude and practices of pesticide use and acetylcholinesterase depression among farm workers in Nepal. Int. J. Environ. Health Res., 22, 401–415.

Badiou, A., Meled, M. & Belzunces, L.P. (2008). Honeybee Apis mellifera acetylcholinesterase—A biomarker to detect deltamethrin exposure. Ecotoxicol. Environ. Saf., 69, 246–253.

Badiou-Bénéteau, A., Benneveau, A., Géret, F., Delatte, H., Becker, N., Brunet, J.L., et al. (2013). Honeybee biomarkers as promising tools to monitor environmental quality. Environ. Int., 60, 31–41.

Belzunces, L.P., Tchamitchian, S. & Brunet, J.-L. (2012). Neural effects of insecticides in the honey bee. Apidologie, 43, 348–370.

Benuszak, J., Laurent, M. & Chauzat, M.-P. (2017). The exposure of honey bees (Apis mellifera; Hymenoptera: Apidae) to pesticides: Room for improvement in research. Sci. Total Environ., 587-588, 423–438.

Boily, M., Sarrasin, B., DeBlois, C., Aras, P. & Chagnon, M. (2013). Acetylcholinesterase in honey bees (Apis mellifera) exposed to neonicotinoids, atrazine and glyphosate: laboratory and field experiments. Environ. Sci. Pollut. Res., 20, 5603–5614.

Buchmann, S.L. & Nabhan, G.P. (1997). The Forgotten Pollinators. Island Press.

Calatayud-Vernich, P., VanEngelsdorp, D. & Picó, Y. (2019). Beeswax cleaning by solvent extraction of pesticides. MethodsX, 6, 980–985.

Chmiel, J.A., Daisley, B.A., Pitek, A.P., Thompson, G.J. & Reid, G. (2020). Understanding the Effects of Sublethal Pesticide Exposure on Honey Bees: A Role for Probiotics as Mediators of Environmental Stress. Front. Ecol. Evol., 8.

Christen, V. & Fent, K. (2017). Exposure of honey bees (Apis mellifera) to different classes of insecticides exhibit distinct molecular effect patterns at concentrations that mimic environmental contamination. Environ. Pollut., 226, 48–59.

Cox-Foster, D.L., Conlan, S., Holmes, E.C., Palacios, G., Evans, J.D., Moran, N.A., et al. (2007). A Metagenomic Survey of Microbes in Honey Bee Colony Collapse Disorder. Science, 318, 283–287.

Ellis, J.D., Evans, J.D. & Pettis, J. (2010). Colony losses, managed colony population decline, and Colony Collapse Disorder in the United States. J. Apic. Res., 49, 134–136.

Ellman, G.L., Courtney, K.D., Andres Jr, V. & Featherstone, R.M. (1961). A new and rapid colorimetric determination of acetylcholinesterase activity. Biochem. Pharmacol., 7, 88–95.

Forfert, N., Troxler, A., Retschnig, G., Gauthier, L., Straub, L., Moritz, R.F.A., et al. (2017). Neonicotinoid pesticides can reduce honeybee colony genetic diversity. PLOS ONE, 12, e0186109.

Frasco, M.F., Fournier, D., Carvalho, F. & Guilhermino, L. (2005). Do metals inhibit acetylcholinesterase (AChE)? Implementation of assay conditions for the use of AChE activity as a biomarker of metal toxicity. Biomarkers, 10, 360–375.

Gallai, N., Salles, J.-M., Settele, J. & Vaissière, B.E. (2009). Economic valuation of the vulnerability of world agriculture confronted with pollinator decline. Ecol. Econ., 68, 810–821.

Gauthier, M., Aras, P., Paquin, J. & Boily, M. (2018). Chronic exposure to imidacloprid or thiamethoxam neonicotinoid causes oxidative damages and alters carotenoid-retinoid levels in caged honey bees (Apis mellifera). Sci. Rep., 8, 16274.

Goulson, D., Nicholls, E., Botías, C. & Rotheray, E.L. (2015). Bee declines driven by combined stress from parasites, pesticides, and lack of flowers. Science, 1255957.

Government of Nepal, National Planning Secretariat. (2014). Population monograph of Nepal. 1st edn. Central Bureau of Statistics.

Hanley, N., Breeze, T.D., Ellis, C. & Goulson, D. (2015). Measuring the economic value of pollination services: Principles, evidence and knowledge gaps. Ecosyst. Serv., 14, 124–132.

Hein, L. (2009). The economic value of the pollination service, a review across scales. Open Ecol. J., 2.

Hung, K.-L.J., Kingston, J.M., Albrecht, M., Holway, D.A. & Kohn, J.R. (2018). The worldwide importance of honey bees as pollinators in natural habitats. Proc. R. Soc. B Biol. Sci., 285, 20172140.

Johnson, R.M., Dahlgren, L., Siegfried, B.D. & Ellis, M.D. (2013). Acaricide, Fungicide and Drug Interactions in Honey Bees (Apis mellifera). PLOS ONE, 8, e54092.

Jones Ritten, C., Peck, D., Ehmke, M. & Patalee, M.A.B. (2018). Firm Efficiency and Returns-to-Scale in the Honey Bee Pollination Services Industry. J. Econ. Entomol., 111, 1014–1022.

Khalifa, M.H., Aly, G.F. & Abdelhameed, K.M.A. (2020). Heavy Metal Accumulation and The Possible Correlation with Acetylcholinesterase Levels in Honey Bees from Polluted Areas of Alexandria, Egypt. Afr. Entomol., 28, 385–393.

Kim, S., Kim, K., Lee, J.H., Han, S.H. & Lee, S.H. (2019). Differential expression of acetylcholinesterase 1 in response to various stress factors in honey bee workers. Sci. Rep., 9, 10342.

Kim, Y.H., Cha, D.J., Jung, J.W., Kwon, H.W. & Lee, S.H. (2012). Molecular and Kinetic Properties of Two Acetylcholinesterases from the Western Honey Bee, Apis mellifera. PLOS ONE, 7, e48838.

Kim, Y.H., Kim, J.H., Kim, K. & Lee, S.H. (2017). Expression of acetylcholinesterase 1 is associated with brood rearing status in the honey bee, Apis mellifera. Sci. Rep., 7, 39864.

Lee, C., Jeong, S., Jung, C. & Burgett, M. (2016). Acute oral toxicity of neonicotinoid insecticides to four species of honey bee, Apis florea, A. cerana, A. mellifera, and A. dorsata. J. Apic., 31, 51–58.

Li, Z., Li, M., He, J., Zhao, X., Chaimanee, V., Huang, W.-F., et al. (2017). Differential physiological effects of neonicotinoid insecticides on honey bees: A comparison between Apis mellifera and Apis cerana. Pestic. Biochem. Physiol., 140, 1–8.

Lundin, O., Rundlöf, M., Smith, H.G., Fries, I. & Bommarco, R. (2015). Neonicotinoid Insecticides and Their Impacts on Bees: A Systematic Review of Research Approaches and Identification of Knowledge Gaps. PLOS ONE, 10, e0136928.

Mahapatra, P.S., Jain, S., Shrestha, S., Senapati, S. & Puppala, S.P. (2018). Ambient endotoxin in PM10 and association with inflammatory activity, air pollutants, and meteorology, in Chitwan, Nepal. Sci. Total Environ., 618, 1331–1342.

Marwick, B. & Krishnamoorthy, K. (2019). cvequality: Tests for the Equality of Coefficients of Variation from Multiple Groups. R software package version 0.1.3. R..

Matsuda, K., Buckingham, S.D., Kleier, D., Rauh, J.J., Grauso, M. & Sattelle, D.B. (2001). Neonicotinoids: insecticides acting on insect nicotinic acetylcholine receptors. Trends Pharmacol. Sci., 22, 573–580.

Michener, C.D., McGinley, R.J. & Danforth, B.N. (1994). The bee genera of North and Central America (Hymenoptera: Apoidea). Smithsonian Institution Press.

Mitchell, E. a. D., Mulhauser, B., Mulot, M., Mutabazi, A., Glauser, G. & Aebi, A. (2017). A worldwide survey of neonicotinoids in honey. Science, 358, 109–111.

Morse, R.A. & Calderone, N.W. (2000). The value of honey bees as pollinators of US crops in 2000. Bee Cult., 128, 1–15.

Neupane, B.B., Sharma, A., Giri, B. & Joshi, M.K. (2020). Characterization of airborne dust samples collected from core areas of Kathmandu Valley. Heliyon, 6, e03791.

Ollerton, J., Price, V., Armbruster, W.S., Memmott, J., Watts, S., Waser, N.M., et al. (2012). Overplaying the role of honey bees as pollinators: a comment on Aebi and Neumann (2011). Trends Ecol. Evol., 27, 141.

Palmer, M.J., Moffat, C., Saranzewa, N., Harvey, J., Wright, G.A. & Connolly, C.N. (2013). Cholinergic pesticides cause mushroom body neuronal inactivation in honeybees. Nat. Commun., 4, 1634.

Pokhrel, S., Thapa, R.B., Neupane, F.P. & Shrestha, S.M. (2006). Absconding Behavior and Management of Apis cerana F. Honeybee in Chitwan, Nepal. J. Inst. Agric. Anim. Sci., 27, 77–86.

Porrini, C., Ghini, S., Girotti, S., Sabatini, A.G., Gattavecchia, E. & Celli, G. (2002). Use of honey bees as bioindicators of environmental pollution in Italy. In: Honey bees: estimating the environmental impact of chemicals (eds. Devillers, J. & Pham-Delegue, M.-H.). Taylor and Francis, London, pp. 186–247.

Potts, S.G., Biesmeijer, J.C., Kremen, C., Neumann, P., Schweiger, O. & Kunin, W.E. (2010). Global pollinator declines: trends, impacts and drivers. Trends Ecol. Evol., 25, 345–353.

Prescott-Allen, R. & Prescott-Allen, C. (1990). How Many Plants Feed the World? Conserv. Biol., 4, 365–374.

Product information: Acetylcholinesterase Activity Assay Kit. (2013)..

Rajarathnam, U., Athalye, V., Ragavan, S., Maithel, S., Lalchandani, D., Kumar, S., et al. (2014). Assessment of air pollutant emissions from brick kilns. Atmos. Environ., 98, 549–553.

Samson-Robert, O., Labrie, G., Mercier, P.-L., Chagnon, M., Derome, N. & Fournier, V. (2015). Increased Acetylcholinesterase Expression in Bumble Bees During Neonicotinoid-Coated Corn Sowing. Sci. Rep., 5, 12636.

Scharlaken, B., Graaf, D.C.D., Memmi, S., Devreese, B., Beeumen, J.V. & Jacobs, F.J. (2007). Differential protein expression in the honey bee head after a bacterial challenge. Arch. Insect Biochem. Physiol., 65, 223–237.

Shrestha, A. (2016). Air pollution levels higher in Bhaktapur than in Kathmandu. Himal. Times. Available at: https://thehimalayantimes.com/kathmandu/air-pollution-levels-higher-bhaktapur-kathmandu/. Last accessed 27 December 2020.

Straub, L., Williams, G.R., Vidondo, B., Khongphinitbunjong, K., Retschnig, G., Schneeberger, A., et al. (2019). Neonicotinoids and ectoparasitic mites synergistically impact honeybees. Sci. Rep., 9, 8159.

Tan, J., Galligan, J.J. & Hollingworth, R.M. (2007). Agonist actions of neonicotinoids on nicotinic acetylcholine receptors expressed by cockroach neurons. NeuroToxicology, 28, 829–842.

Tan, K., Chen, W., Dong, S., Liu, X., Wang, Y. & Nieh, J.C. (2015). A neonicotinoid impairs olfactory learning in Asian honey bees (Apis cerana) exposed as larvae or as adults. Sci. Rep., 5, 10989.

Thapa, R., Aryal, S. & Jung, C. (2018). Beekeeping and Honey Hunting in Nepal: Current Status and Future Perspectives. In: Asian Beekeeping in the 21st Century (eds. Chantawannakul, P., Williams, G. & Neumann, P.). Springer Singapore, Singapore, pp. 111–127.

Vanbergen, A.J. & Initiative, the I.P. (2013). Threats to an ecosystem service: pressures on pollinators. Front. Ecol. Environ., 11, 251–259.

vanEngelsdorp, D. & Meixner, M.D. (2010). A historical review of managed honey bee populations in Europe and the United States and the factors that may affect them. J. Invertebr. Pathol., 103, Supplement, S80–S95.

Walker, K. (2006). Common Honeybee (Apis mellifera). Available at: https://www.padil.gov.au/pests-and-diseases/pest/main/135540/8765. Last accessed 31 December 2020.

Weinstock, G.M., Robinson, G.E., Gibbs, R.A., Worley, K.C., Evans, J.D., Maleszka, R., et al. (2006). Insights into social insects from the genome of the honeybee Apis mellifera. Nature, 443, 931–949.

Williamson, S.M., Moffat, C., Gomersall, M., Saranzewa, N., Connolly, C. & Wright, G.A. (2013). Exposure to Acetylcholinesterase Inhibitors Alters the Physiology and Motor Function of Honeybees. Front. Physiol. 4.

Wu-Smart, J. & Spivak, M. (2016). Sub-lethal effects of dietary neonicotinoid insecticide exposure on honey bee queen fecundity and colony development. Sci. Rep., 6, 32108.

